# A Network-Based Approach for Prioritizing Candidate Genes in Alzheimer’s Disease

**DOI:** 10.64898/2026.03.10.710766

**Authors:** Nakshatra Malhotra, Sibendu Samanta, Anuj Deshpande

## Abstract

Alzheimer’s disease (AD) is a multifactorial neurodegenerative disorder characterized by coordinated dysregulation of multiple genes, requiring system-level approaches to elucidate underlying molecular mechanisms. While existing computational studies largely focus on differential expression analysis or machine–learning–based feature selection, they often overlook inter-gene relationships and network topology, limiting biological interpretability. In this study, we present a network-based framework for prioritizing candidate genes in Alzheimer’s disease by integrating gene co-expression network analysis with multiple centrality measures. Transcriptomic data comprising approximately 39,000 genes across 324 Alzheimer’s and control samples were preprocessed using log-transformation, variance filtering and Z-score normalization, followed by LASSO-based feature selection to retain phenotype-associated genes. A weighted gene co-expression network was constructed using Pearson correlation to capture coordinated transcriptional activity. Network topology was analyzed using degree, betweenness and eigenvector centrality to identify genes that are highly connected, act as information brokers or occupy influential positions within the network. A consensus ranking was derived by merging these centrality measures, enabling robust prioritization of candidate genes. The results highlight a subset of highly central genes, including several small nucleolar RNAs and regulatory transcripts implicated in RNA processing, synaptic function and neurodegenerative pathways. By jointly leveraging co-expression structure and complementary centrality metrics, the proposed framework provides an interpretable and reproducible strategy for identifying biologically meaningful gene candidates, offering potential utility for biomarker discovery and therapeutic target prioritization in Alzheimer’s disease.

## I. INTRODUCTION

**A**LZHEIMER’S Alzheimer’s disease (AD) is the most prevalent neurodegenerative disorder globally, accounting for nearly 60-70% of dementia cases among the elderly population. According to the World Health Organization (2024), more than 55 million individuals currently live with dementia worldwide and approximately 10 million new cases are reported annually [1]. In India alone, epidemiological analyses suggest that more than 8 million individuals are currently affected, with projections indicating a threefold increase by 2050 [2]. Despite decades of extensive molecular, pathological and clinical research, the underlying mechanisms of AD remain incompletely understood due to its polygenic, heterogeneous and multifactorial nature [3]. The disease encompasses a complex interplay between amyloid processing, tau aggregation, synaptic dysfunction, chronic inflammation, mitochondrial impairment and genetic susceptibility. Traditional diagnostic procedures rely on cognitive assessment, neuro-imaging modalities (MRI, PET) and invasive cerebrospinal fluid (CSF) biomarker assays such as amyloid-*β* (A*β*42), total tau, and phosphorylated tau [4]. Although these approaches remain clinically valuable, they are often costly, invasive and insufficiently sensitive for detecting the earliest stages of AD—long before severe neuronal loss occurs. Therapeutic strategies such as cholinesterase inhibitors and NMDA receptor antagonists provide symptomatic relief but do not halt disease progression [5]. Consequently, the need for early, accurate and biologically interpretable diagnostic and predictive markers has never been greater.

A growing body of evidence demonstrates that genetic factors shape susceptibility, progression and molecular sub-types of AD. Genome-wide association studies (GWAS), RNA-sequencing (RNA-Seq) and pathway enrichment analyses have identified major risk loci including *APOE ε*4, *TREM2, CLU* and *PICALM* [7], [8]. However, although these approaches highlight important individual genes, they frequently lack reproducibility and often fail to capture the underlying co-regulatory behavior that emerges only when genes are studied collectively. AD pathogenesis is inherently network-driven; isolated markers rarely explain the full spectrum of molecular dysfunction and single-gene studies fail to capture interdependencies critical to neuronal health [6].

Contemporary computational approaches, especially machine learning (ML)-based classification and feature selection, have accelerated AD biomarker discovery. Techniques such as LASSO, support vector machines, random forests, gradient boosting and AutoML frameworks demonstrate strong predictive performance. Yet, traditional ML pipelines typically treat genes as independent features, disregarding their biological relationships and regulatory hierarchies [9]. This results in feature sets that are predictive but not necessarily mechanistically meaningful, limiting downstream interpretability and biological validation.

To overcome these challenges, research has increasingly shifted toward genetic network analysis—specifically, gene co-expression networks (GCNs), gene regulatory networks (GRNs) and hybrid graph-based models. These approaches capture correlated or causal relationships between genes, revealing functional modules associated with amyloid metabolism, tau phosphorylation, synaptic signaling, RNA processing, oxidative stress and neuroinflammation [10]– [12]. Unlike single-gene analyses, network-based frameworks describe the broader molecular landscape and identify gene communities or hubs that may act as master regulators or central coordinators of disease-specific pathways.

However, network-only methods introduce their own limitations. Co-expression measures are sensitive to thresholding, batch effects and noise. Most studies rely on a single centrality metric (e.g., degree), which may oversimplify node importance. Moreover, few existing frameworks integrate predictive ML with network topology, resulting in hub genes that are central in topology but not necessarily relevant to phenotype discrimination. Hence, a key research need is to combine the predictive strength of ML with the biological interpretability of network-based centrality in a systematic, reproducible and data-driven manner. To address these gaps, we introduce an automated, modular computational pipeline that integrates—LASSO-based feature selection to identify genes strongly associated with AD phenotype; AutoML optimization (TPOT) and classical supervised models to ensure robust, reproducible classification performance; Weighted gene co-expression network analysis (GCN) to construct biologically meaningful interaction maps; Multi-metric network centrality analysis (degree, betweenness, eigenvector) to prioritize key hubs; Consensus centrality scoring to generate a biologically interpretable ranked gene list.

This integrative framework emphasizes both prediction accuracy and biological insight. Unlike conventional DEG or ML-only approaches, our pipeline operates at a systems level, characterizing molecular dysregulation through interconnected gene modules rather than isolated markers. By combining robust feature-selection with network topology, A-DANCE identifies regulatory hubs that are both discriminative and structurally important, addressing long-standing issues of instability, non-reproducibility and lack of biological interpretability.

Gene co-expression networks (GCNs) capture coordinated transcriptional activity, allowing researchers to identify groups of genes that act together in biological pathways [10], [11]. In AD, such network models have proven essential for uncovering molecular signatures linked to amyloid metabolism, tau aggregation, mitochondrial dysfunction, RNA splicing and microglial activation [12]. By mapping correlations between thousands of genes across affected and healthy subjects, GCNs reveal structure that would be invisible to conventional feature-by-feature analyses.

In the present study, we analyzed 39,000 genes across 324 AD and control samples. After normalization and preprocessing (log_2_ transformation, variance filtering, Z-score standardization), LASSO regression identified 280 genes with strong predictive relevance. Supervised ML models—including Logistic Regression, Random Forest, XG-Boost, and TPOT AutoML—were trained to evaluate classification performance. Logistic Regression achieved the highest accuracy (0.875) with a ROC AUC of 0.9474, indicating that selected gene signatures provide substantial discriminatory power. Weighted co-expression networks were subsequently constructed using Pearson correlation thresholds (r > 0.70). Louvain community detection revealed tightly connected gene clusters. Centrality metrics were applied to identify influential network hubs, with small nucleolar RNAs (e.g., *SNORD115-6, SNORD116-9, SNORA63D*) emerging as key regulators-consistent with recent studies implicating RNA-processing pathways in neuro-degeneration.

## II. METHODOLOGY

### A. GENETIC NETWORK FOR ALZHEIMER’S DISEASE

Starting from normalized expression (FPKM → log_2_(*x* + 1), optionally TPM), we apply variance filtering (retain top N genes by variance-e.g., top 2,000) to reduce noise. Pairwise Pearson correlation (*r*_*ij*_) is computed between genes (i) and (j) across samples; an adjacency matrix (A) is defined as a weighted matrix with entries:

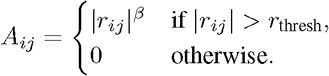

where *r*_thresh_ is a chosen correlation threshold (e.g., 0.7) and *β* is an optional soft power (as used in WGCNA) to emphasize strong correlations [10]. The resulting weighted undirected graph *G* = (*V, E*) has nodes *V* (genes) and weighted edges *E* (co-expression links). Community detection (Louvain) can be applied to *G* to extract modules. Empirically, this pipeline has revealed modules enriched for synaptic, mitochondrial or RNA-processing pathways in AD cohorts.

**FIGURE 1.**
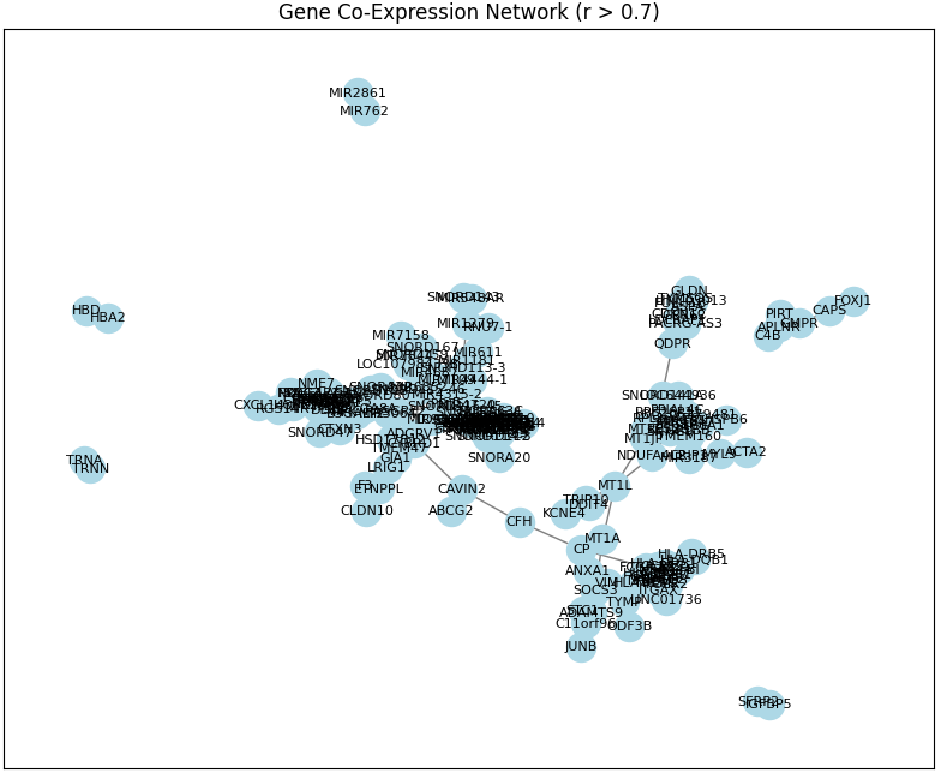
?

**FIGURE 2.**
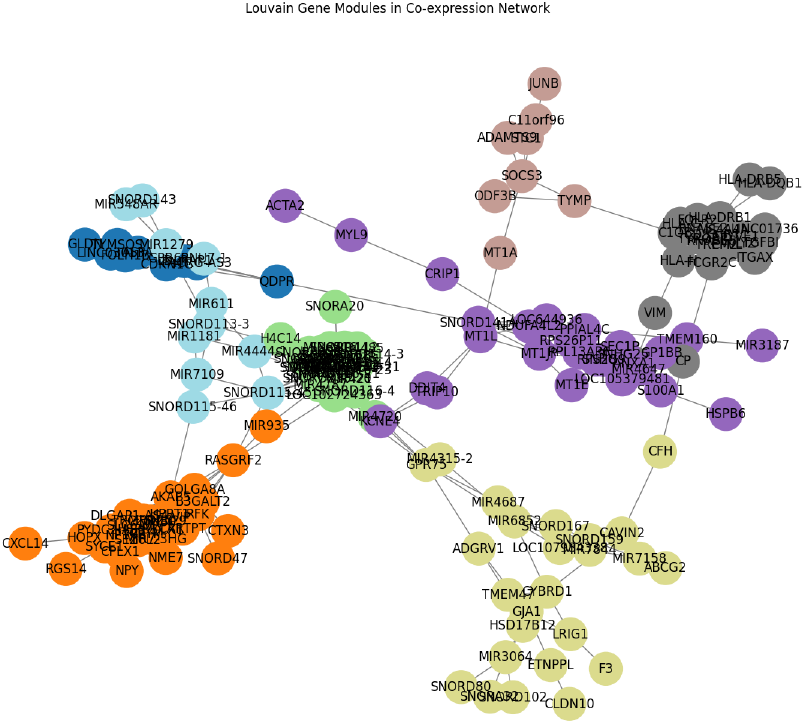
?

### B. CENTRALITY

We use three complementary centrality metrics—degree centrality, betweenness centrality, and eigenvector centrality—and combine them into a consensus ranking. These measures capture complementary aspects of node importance in complex networks and are widely used in graph theory, network science, and biological network analysis [14]–[16], [18].

#### 1) Degree Centrality (weighted)

For a weighted graph *G* with adjacency matrix *A*, the weighted degree (sometimes called node strength) of node *i* is defined as

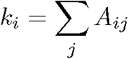

Here, *i* denotes a focal gene in the network and *j* indexes all other genes directly connected to gene *i*. The weighted adjacency matrix element *A*_*ij*_ represents the strength of the functional association between genes *i* and *j*, such as co-expression correlation, regulatory influence, or protein– protein interaction confidence [16], [18]. The summation over *j* therefore aggregates the total interaction strength of gene *i* with its immediate neighbors. The quantity *k*_*i*_ corresponds to the weighted degree (node strength) of gene *i*, reflecting its local connectivity in the gene interaction network [14].

Normalized degree centrality (to range [0, 1]):

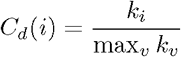

In this expression, *C*_*d*_(*i*) denotes the normalized degree centrality of gene *i*, while max_*v*_ *k*_*v*_ represents the maximum weighted degree observed among all genes *v* in the network. This normalization rescales degree centrality values to the unit interval, allowing direct comparison of local connectivity across genes within the same network [16].

Nodes with high *k*_*i*_ are directly connected to many others or to a few highly weighted partners, suggesting locally influential genes in the interaction network.

#### 2) Betweenness Centrality (shortest-path based)

Betweenness centrality measures the fraction of shortest paths that pass through node *i* [14]. For weighted graphs we compute shortest paths using edge weights converted to lengths (for example, *l*_*ij*_ = 1*/A*_*ij*_ for positive *A*_*ij*_):

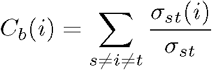

In this formulation, *s* and *t* denote distinct source and target genes in the network, excluding the focal gene *i*. The term *σ*_*st*_ represents the total number of shortest paths between genes *s* and *t*, while *σ*_*st*_(*i*) denotes the number of those shortest paths that pass through gene *i*. Betweenness centrality thus quantifies how frequently gene *i* lies on shortest communication paths connecting other gene pairs in the network [14], [16].

Normalization is applied to allow comparison across networks:

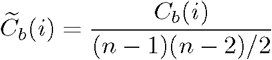

Here, 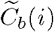 denotes the normalized betweenness centrality of gene *i*, and *n* represents the total number of genes in the network. The normalization factor corresponds to the maximum possible number of gene pairs in an undirected network, ensuring comparability of betweenness values across networks of different sizes [16].

Interpretation: nodes with high *C*_*b*_ control information flow between modules and may represent regulatory bottlenecks or signaling mediators within biological networks [18].

#### 3) Eigenvector Centrality

Eigenvector centrality assigns relative scores to nodes based on their connections to other highly connected nodes [15]. It is defined as the leading eigenvector of the adjacency matrix *A*:

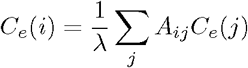

In this equation, *C*_*e*_(*i*) denotes the eigenvector centrality of gene *i, A*_*ij*_ represents the interaction strength between genes *i* and *j*, and *C*_*e*_(*j*) denotes the eigenvector centrality of neighboring gene *j*. The scalar *λ* corresponds to the largest eigenvalue of the adjacency matrix *A* and serves as a normalization constant [15], [16]. Eigenvector centrality assigns higher scores to genes that are connected to other highly influential genes in the network. Thus, a high *C*_*e*_(*i*) implies that gene *i* participates in a cluster of influential or hub-like genes.

#### 4) Combining Centralities (Consensus Score)

Given normalized metrics *Ĉ*_*d*_(*i*), *Ĉ*_*b*_(*i*), and *Ĉ*_*e*_(*i*) (each scaled to [0, 1]), we compute a weighted consensus score:

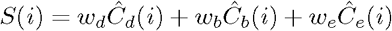

where *w*_*d*_ + *w*_*b*_ + *w*_*e*_ = 1. A reasonable default is equal weighting (*w*_*d*_ = *w*_*b*_ = *w*_*e*_ = 1/3). Alternatively, weights can be learned using a simple logistic regression model on an independent validation set, if known marker genes are available as positive labels [17].

The combined ranking defined by *S*(*i*) produces a robust prioritized gene list. Studies in network biology and systems medicine suggest that combining multiple centrality measures often improves identification of biologically relevant hub genes compared with using a single metric alone [18], [22].

### C. APPLICATION OF CENTRALITY TO GENETIC NETWORK ANALYSIS

To obtain a robust ranking of candidate genes, three complementary network centrality measures—degree centrality, betweenness centrality, and eigenvector centrality—were integrated into a single consensus score. Each measure captures a distinct topological role of a gene within the co-expression network: degree centrality reflects local connectivity, betweenness centrality quantifies a gene’s role in mediating information flow between network modules, and eigenvector centrality measures global influence by accounting for connections to highly connected neighbors [14]–[16], [18].

For each gene *i*, raw centrality values were first normalized using min–max scaling to ensure comparability across different metrics [23]:

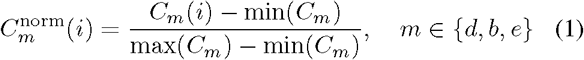

Here, *C*_*m*_(*i*) denotes the raw centrality value of gene *i* for metric *m*, where *m* ∈ {*d, b, e}* corresponds to degree, betweenness, and eigenvector centrality, respectively. The terms min(*C*_*m*_) and max(*C*_*m*_) represent the minimum and maximum values of centrality *m* across all genes in the network. This transformation rescales each centrality measure to a common range, enabling integration of multiple centrality metrics.

The normalized centrality measures were then combined using an unweighted linear aggregation to compute a consensus centrality score:

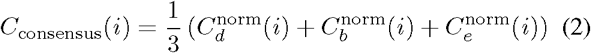

In this expression, *C*_consensus_(*i*) denotes the consensus centrality score of gene *i*, while 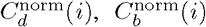, and 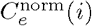 represent the normalized degree, betweenness, and eigenvector centrality values of gene *i*, respectively. Equal weighting of the three components reflects the assumption that local connectivity, global mediation, and hierarchical influence contribute comparably to overall gene importance.

Genes were ranked in descending order of *C*_consensus_(*i*), with higher values indicating greater overall topological importance within the gene co-expression network. This consensus-based approach integrates local, intermediary, and global network characteristics, thereby reducing metric-specific bias and yielding a stable and biologically interpretable prioritization of Alzheimer’s disease–associated candidate genes [18], [22].

**FIGURE 3.**
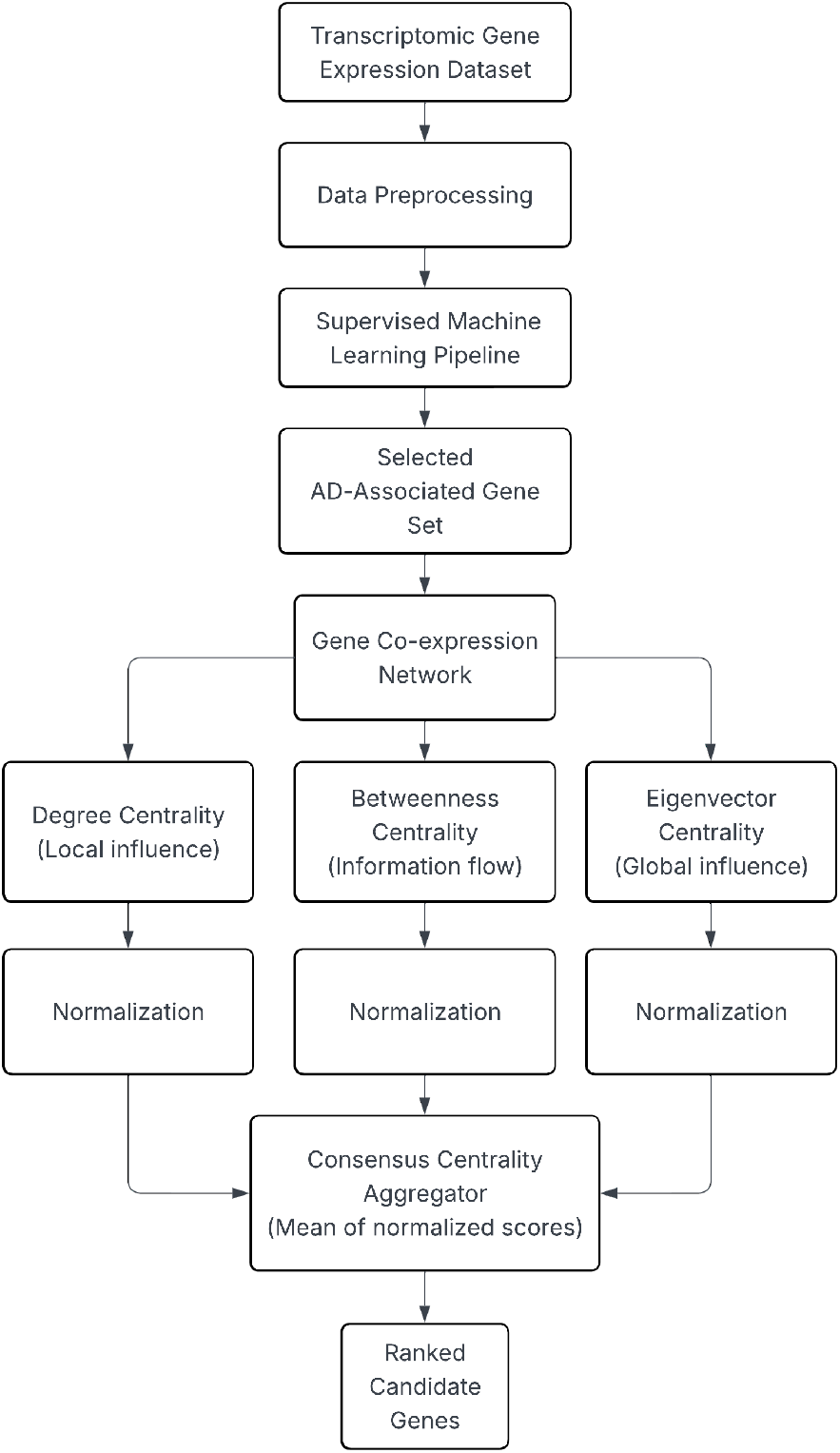
Workflow for the machine learning and network analysis pipeline used to identify candidate Alzheimer’s disease genes.

## III. RESULTS

**FIGURE 4.**
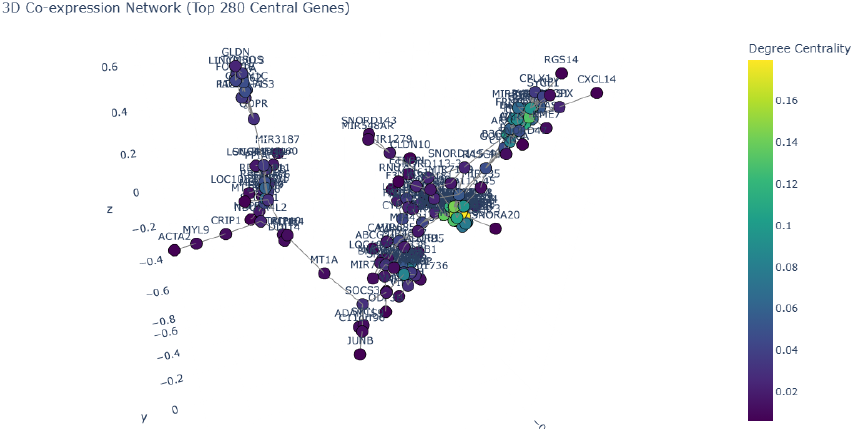
Degree centrality network showing locally connected hub genes.

The proposed network-based framework was applied to transcriptomic data comprising 324 Alzheimer’s disease (AD) and control samples. Following normalization and variance-based filtering, a reduced gene set was used to construct a weighted gene co-expression network using Pearson correlation thresholds. Gene co-expression networks provide a powerful framework for identifying coordinated transcriptional activity and functional gene modules [10], [11].

**FIGURE 5.**
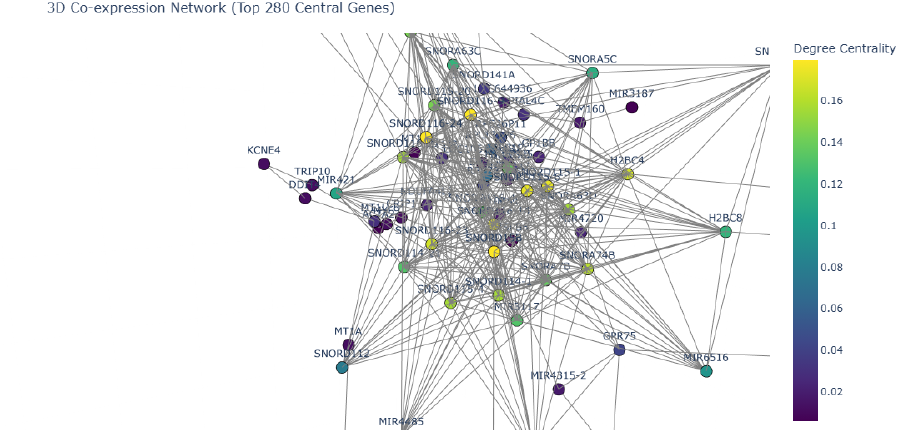
Betweenness centrality network highlighting genes acting as bridges between functional modules.

**FIGURE 6.**
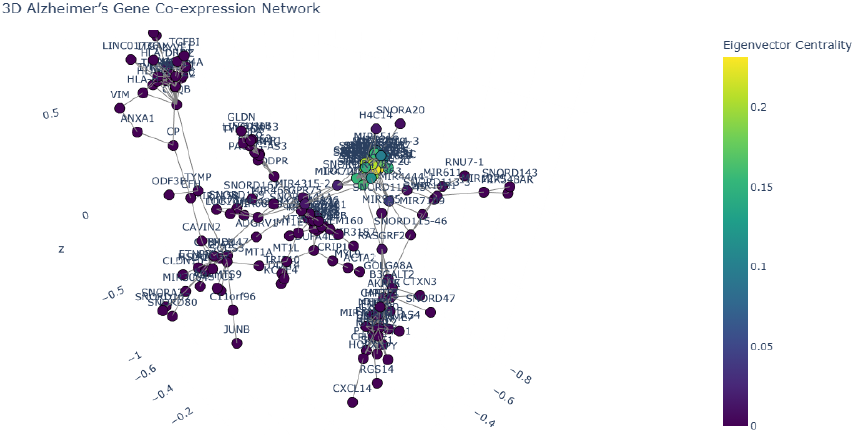
Eigenvector centrality network illustrating globally influential genes connected to other highly connected nodes.

**FIGURE 7.**
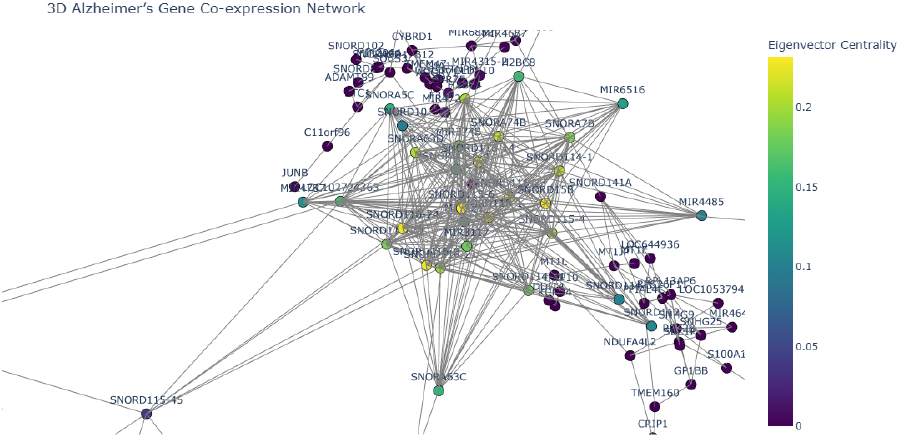
Eigenvector centrality distribution across the gene interaction network.

The resulting network exhibited a modular topology with a dominant connected component, indicating coordinated transcriptional activity among disease-associated genes. Such modular organization is commonly observed in biological systems and reflects the functional architecture of cellular processes [18], [24].

Centrality analysis was performed using degree, betweenness, and eigenvector measures to quantify gene importance from complementary topological perspectives. After normalization, the three metrics were integrated into a consensus centrality score, enabling robust prioritization of candidate genes.

Genes ranked at the top of the consensus list demonstrated high connectivity, strong intermediary roles between modules, and association with globally influential network regions. Previous studies have shown that highly central genes in biological networks often correspond to essential regulatory components and disease-associated hubs [22].

Functional inspection of highly ranked genes revealed enrichment in biological processes related to RNA processing, synaptic signaling, and neurodegenerative pathways. Notably, several small nucleolar RNAs and transcription-related genes emerged as central hubs, suggesting a potential regulatory role in Alzheimer’s disease progression.

These findings indicate that network topology–driven prioritization captures biologically meaningful gene candidates beyond those identified through differential expression analysis alone.

The complete ranked list of genes and their corresponding centrality scores are provided in the Supplementary Material.

## IV. DISCUSSION

This study demonstrates that integrating gene co-expression networks with centrality-based ranking provides valuable insights into the molecular organization underlying Alzheimer’s disease. Unlike conventional approaches that focus on isolated genes or purely predictive machine learning features, the proposed framework emphasizes system-level interactions and network structure, which are critical for understanding complex neurodegenerative disorders [10], [18], [24].

The identification of highly central genes involved in RNA metabolism and synaptic function aligns with emerging evidence that post-transcriptional regulation and neuronal communication pathways are disrupted early in Alzheimer’s disease [3], [6], [7]. Importantly, genes with high degree centrality reflect strong co-regulatory influence, while high betweenness centrality highlights genes that mediate information flow between functional modules. Eigenvector centrality further prioritizes genes embedded within globally influential regions of the network. Their integration ensures that prioritized genes are not only locally connected but also structurally and globally significant [14]–[16].

Compared with previous genetic studies relying solely on genome-wide association studies (GWAS) or differential expression analysis, the network-based approach reduces sensitivity to dataset-specific noise and improves biological interpretability [7], [8]. Although the method is influenced by correlation thresholds and network construction parameters, the consensus centrality strategy mitigates these effects by emphasizing stability across multiple metrics. Future work incorporating multi-omic data and longitudinal cohorts may further strengthen the translational relevance of the identified candidates.

A limitation of the current framework is that gene co-expression relationships inferred from bulk transcriptomic data may not fully capture cell-type specific regulatory interactions in the brain. Future studies integrating single-cell transcriptomics and spatial transcriptomics may provide a more refined view of disease-associated regulatory networks.

## V. CONCLUSION

In this work, we present a network-based framework for prioritizing candidate genes in Alzheimer’s disease by integrating gene co-expression analysis with multi-metric centrality scoring. By combining degree, betweenness and eigenvector centralities into a consensus ranking, the proposed approach identifies genes that are topologically influential across multiple levels of network organization. The results highlight biologically relevant gene candidates associated with RNA processing and neurodegenerative pathways, demonstrating the effectiveness of centrality-driven network analysis for uncovering system-level disease mechanisms. This framework offers a transparent, scalable and biologically interpretable strategy for gene prioritization and may serve as a foundation for future biomarker discovery and therapeutic target identification in Alzheimer’s disease.

